# Heat and Nitrate Drive Metabolic and Immune Reprogramming Leading to the Collapse of Symbiosis in the Model Sea Anemone Aiptasia

**DOI:** 10.64898/2026.05.19.726363

**Authors:** Jeric Da-Anoy, Benjamin Glass, Oliwia Jasnos, Kyle S. Toyoma, Chloe Costa, Kian Thompson, Heather Donnelly, Mu-Han Chen, Federica Calabrese, Lorenz Ponce, Maria Valadez-Ingersoll, Kamel Ramzi Moufarrej, Aden Nagree, Alex Geisser, Maria Medalla, Jeffrey Marlow, Robinson W. Fulweiler, Xingchen Wang, Thomas D. Gilmore, Sarah W. Davies

**Affiliations:** Department of Biology, Boston University, Boston, MA, USA; Department of Earth and Environmental Sciences, Boston College, Newton, MA, USA; Department of Biology, Woods Hole Oceanographic Institution, Woods Hole, MA, USA; Department of Anatomy and Neurobiology, Boston University School of Medicine, MA, USA

## Abstract

The maintenance of endosymbiosis in cnidarians depends on the tight regulation of host immunity, cell cycle, and nutrient exchange, yet how these processes are impacted by interacting environmental stressors remains largely unknown. To address this, we employed physiological metrics, gene expression analysis, microbiome characterization, imaging (NF-κB localization, endoplasmic reticulum ultrastructure, EdU labeling), and stable isotope tracing in the model sea anemone *Exaiptasia diaphana* to examine the effects of heat and nitrate on these regulatory processes, individually and in combination. Heat treatment led to NF-κB activation, proteostatic stress, suppression of nutrient exchange, decreased cell-cycle progression, and microbiome restructuring, with all effects more pronounced in symbiotic than aposymbiotic anemones. In symbiotic anemones, nitrate partially offset these heat-induced responses through sustained carbon translocation, suggesting that the presence of symbionts, in conjunction with elevated nitrate, can temporarily buffer host thermal stress. However, prolonged combined exposure resulted in holobiont failure. These findings reveal that while nitrate enrichment can transiently delay the onset of bleaching, it does not preserve the regulatory networks required for symbiotic stability — underscoring the vulnerability of cnidarian holobionts to the compounding effects of warming and nitrate pollution.

## INTRODUCTION

Endosymbiosis between reef-building corals and their dinoflagellate algal symbionts (family Symbiodiniaceae) underpins the integrity of coral reef ecosystems^1^. Along with their associated microbial communities, the coral host and its algal symbionts together constitute the cnidarian holobiont. In this association, algal symbionts are maintained within coral host gastrodermal cells inside a specialized, host membrane-derived compartment termed the symbiosome^2^. This symbiosis is primarily nutritional, with coral hosts meeting the majority of their energetic demands through photosynthates produced by symbionts, while the host provides metabolic byproducts including CO_2_ and nitrogenous compounds^3^. This mutualism has enabled corals to thrive in oligotrophic environments^4^, contributing to their evolutionary success^1,5^. Symbiosis stability and, by extension, coral persistence depend on host regulation of the cellular and metabolic processes that maintain symbiosis^3,6^.

In its homeostatic state, coral-algal symbiosis is actively maintained by the host through regulation of immunity^7^, the cell cycle^8^, and nutrient exchange across the symbiosome membrane^9^. Central to both initiation and long-term persistence of this partnership is suppression of host innate immunity in order to tolerate symbionts^7^. For example, decreased expression of the innate immune transcription factor nuclear factor-κB (NF-κB) has been observed during the onset of symbiosis^10,11^. Further, exposure to stressors (e.g., heat) can lead to the reactivation of NF-κB coincident with symbiont loss (i.e., bleaching)^7,10^. Yet, the molecular mechanisms by which stressors influence NF-κB signaling and downstream effects on symbiosis remain poorly characterized^12^. Following the establishment of symbiosis, the host exerts stringent control over symbiont population density to prevent overgrowth^13^. Hosts can control symbiont cell density through interference with symbiont G_2_/M mitotic checkpoint progression^3^, and by imposing nitrogen limitation^14^ that restricts symbiont growth while encouraging continued translocation of fixed carbon to the host. Together, immunity, cell-cycle regulation, and nutrient exchange processes serve as fundamental biological processes required for symbiosis maintenance.

Increasing seawater temperatures threaten coral-algal symbioses on a global scale^15^. Thermal stress disrupts the host’s regulatory control and destabilizes the symbiosis^16^, resulting in dysbiosis and bleaching (i.e., symbiont loss)^6^. One proposed mechanism that initiates dysbiosis is heat-induced impairment of algal photosystems^17^, which leads to excessive production of reactive oxygen species (ROS)^18^. Oxidative stress leads to diffusion of oxidants into the host cytoplasm, triggering expulsion and/or degradation of symbionts^19^. Oxidative stress is also accompanied by proteostatic and endoplasmic reticulum (ER) stress resulting from heat-induced unfolding of proteins and membrane instability^20^. In response to proteostatic and oxidative stress, the host can upregulate molecular pathways associated with stress responses, immunity, and metabolism^21–23^. Heat stress can also disrupt symbiotic nutrient exchange^24^, leaving the host in a carbon-limited state^9^ that can lead to mortality under prolonged or severe stress.

Local stressors, including nutrient enrichment, have been linked to enhanced coral bleaching and disease outbreaks^25^. For example, excess nitrogen has been shown to disrupt host control of algal symbiont density by skewing the holobiont carbon-to-nitrogen (C:N) balance^26^. This shift in the C:N balance disrupts the symbiont nitrogen limitation that normally favors efficient carbon translocation to the host^9^. Increased nutrient (e.g., nitrate) levels can also interact with heat stress to disrupt host physiology and the control of bacterial microbiome communities^27^. Notably, such negative effects can occur even in the absence of bleaching, emphasizing that, although bleaching is a clear visual indicator of dysbiosis^28^, it captures only the final stage of symbiosis breakdown. Indeed, abiotic stressors can perturb carbon metabolism^9^ and calcification^29^ in stony corals even in the absence of bleaching, suggesting that dysbiosis begins before visible algal loss. Despite growing awareness that heat and nutrient enrichment contribute to coral decline, the mechanisms by which these two stressors work in combination to disrupt the core regulatory processes that maintain symbiosis remain largely unknown.

Here, we have used the cnidarian-algal symbiosis model *Exaiptasia diaphana* (hereafter “Aiptasia”)—a symbiotic sea anemone in phylum Cnidaria with corals—to explore how heat and nitrate enrichment reshape host and symbiont physiology, metabolism, gene expression, and bacterial microbiome composition. Specifically, we have examined how these stressors influence three core processes implicated in symbiosis maintenance: host immune regulation, cell-cycle control, and nutrient exchange. Aiptasia is a useful organism in which to address these processes because these anemones can exist for prolonged periods without symbionts (i.e., aposymbiotic), permitting characterization of how stressors impact host outcomes both directly, through effects on host physiology, and indirectly, through effects on symbiosis. By comparing the responses of symbiotic and aposymbiotic Aiptasia to heat, nitrate, and their combination, we have characterized how the presence of algal symbionts alters host stress responses, and how nitrate influences the trajectory of symbiosis breakdown under thermal stress. Through integration of physiological, cellular, and molecular approaches, this work investigates the core mechanisms that sustain the cnidarian-algal mutualism, providing insights into how global and local stressors interact to disrupt this vital symbiosis.

## RESULTS

### Symbiosis promotes physiological collapse in Aiptasia under heat and nitrate stress

To determine how symbiotic state affects stress responses of the Aiptasia holobiont, we exposed aposymbiotic and symbiotic Aiptasia of three genetic strains (CC7, H2, and VWB) to nitrate (5 µM), heat (ramp 1°C per day for 10 days to 34°C), and combined heat + nitrate (Fig. 1A). Under heat, symbiotic Aiptasia showed morphological and physiological signs of stress, including contracted body columns and shortened tentacles (Fig. 1B). Heat also led to deterioration of algal symbiont health, as evidenced by disrupted symbiont cell ultrastructure (e.g., irregular shape, disrupted chloroplast structure, presence of crystalline aggregates^30^, Fig. 1C–D). These morphological and ultrastructural changes in response to heat were accompanied by several physiological responses. Total protein content was significantly reduced in animals exposed to heat compared to controls (25.0 ± 2.6% decrease, t(48) = 7.644, p < 0.0001, Fig. 1E, table S1), whereas carbohydrate content showed no significant treatment effects (F(3,48) = 0.116, p = 0.951, Fig. 1F, table S2). Immune enzyme activity also increased significantly under heat (peroxidase: 69.7 ± 18.3% increase, t(48) = 9.065, p < 0.0001, Fig. 1G, table S3, phenoloxidase: 48.4 ± 8.0% increase, t(48) = 7.134, p < 0.0001, Fig. 1H, table S4). Symbiont photophysiology and density, as evidenced by reduced photochemical efficiency (F_v_/F_m_, 31.3 ± 3.2% decrease, t(120) = 9.909, p < 0.0001, Fig. 1I, table S5) and cell density (27.5 ± 2.5% decrease, t(48) = 10.806, p < 0.0001, Fig. 1J, table S6), respectively, were also impaired by heat treatment. Similar to heat, nitrate enrichment alone led to significantly reduced host total protein content (16.4 ± 2.7% decrease, t(48) = 4.766, p = 0.0001, Fig. 1E, table S1), increased peroxidase activity (73.2 ± 12.7% increase, t(48) = 6.519, p < 0.0001, Fig. 1G, table S3), and reduced algal symbiont density (12.3 ± 2.5% decrease, t(48) = 4.835, p = 0.0001, Fig. 1I, table S6).

**Fig 1.**
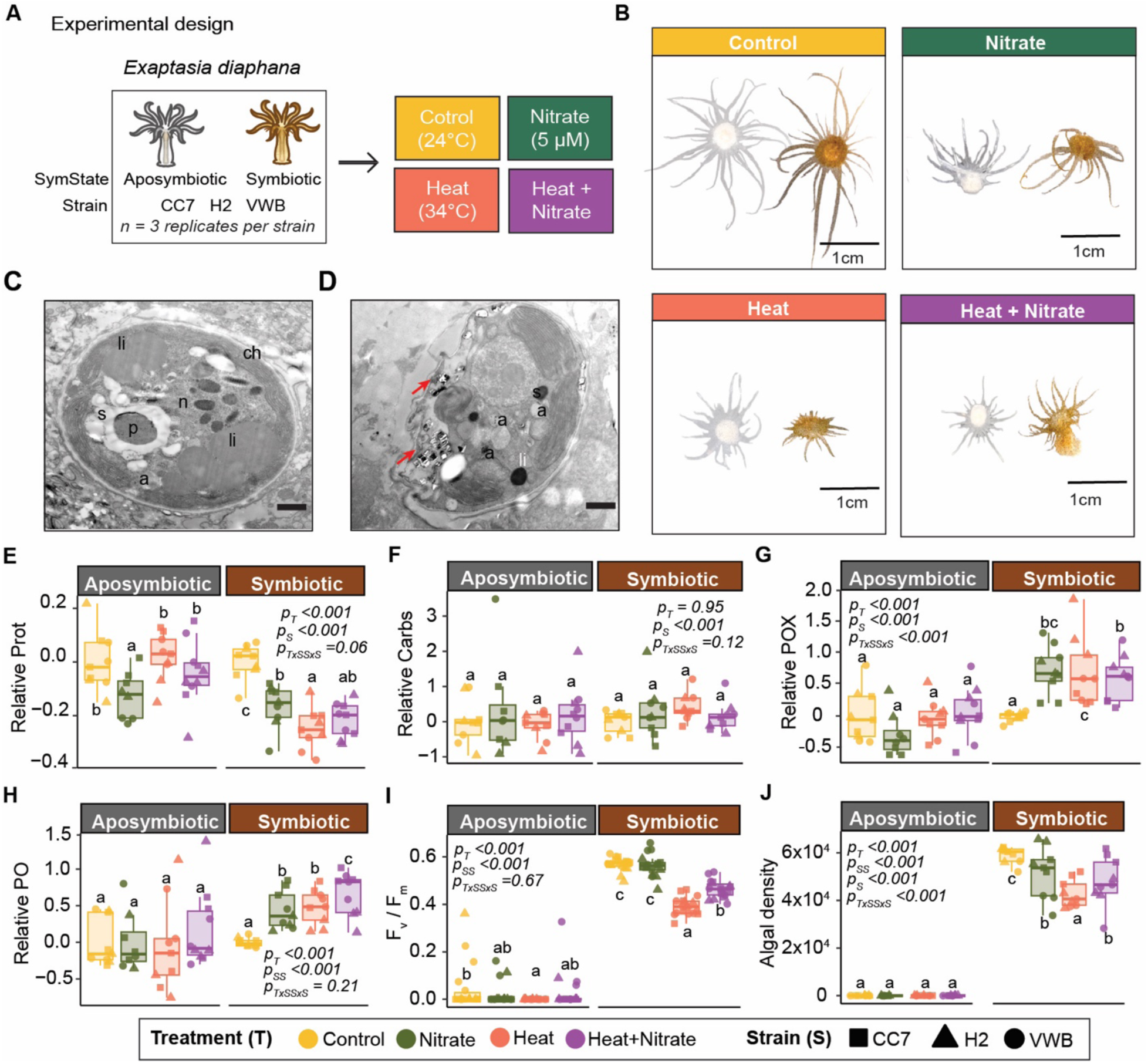
Aiptasia host and algal symbiont physiological responses to heat and nitrate treatments. (A) Experimental design of the study. Symbiotic and aposymbiotic Aiptasia of three genetic strains were exposed to heat, nitrate, or their combination, and responses were compared to control animals not exposed to stress. (B) Representative photos of morphological phenotypes of Aiptasia across treatments at experimental day 12. (C–D) Transmission electron microscopy (TEM) micrographs of (C) intact and (D) damaged algal symbionts within host tissues. Damaged algal symbionts within host tissues exhibited irregular shapes, disrupted chloroplast membranes, more diffuse chromatin within nuclei, and presence of crystalline aggregates (red arrows; uric acid^30,31^). (E) Relative host total protein and (F) carbohydrate content, normalized to wet weight (N = 9 animals per symbiotic state per treatment). (G) Peroxidase (POX) and (H) phenoloxidase (PO) activity, normalized to host protein (N = 9 animals per symbiotic state per treatment). (I) Maximum photochemical efficiency (F_v_/F_m_) and (J) algal cell density across treatments. Values in (E–J) are relative to the control average by symbiotic state. Each point represents one individual. Colors represent treatments (yellow = control, green = nitrate, orange = heat, purple = heat + nitrate); panel header colors denote symbiotic state (grey = Aposymbiotic, brown = Symbiotic); shapes indicate strains (square = CC7, triangle = H2, circle = VWB). ANOVA p-values: P_T_ = treatment, P_S_ = strain, P_SS_ = symbiotic state, P_TxSS_ = interaction.

Under combined heat and nitrate (“heat + nitrate”), symbiotic animals showed less severe morphological declines compared to those exposed to heat alone (Fig. 1B), with only 22.2% developing stress phenotypes compared to 37.0% under heat alone (χ^2^ = 3.93, df = 1, p = 0.047, Fig. 1B, fig. S1). Despite reduced morphological decline under heat + nitrate, anemones exposed to these conditions still showed a stress response including reduced total protein (21.8 ± 2.2% decrease, t(48) = 6.465, p < 0.0001, Fig. 1E, table S1), increased immune enzyme activity (peroxidase: 60.8 ± 11.6% increase, t(48) = 6.222, p < 0.0001, Fig. 1G, table S3; phenoloxidase: 66.3 ± 9.9% increase, t(48) = 9.885, p < 0.0001, Fig. 1H, table S4), decreased F_v_/F_m_ (18.6 ± 3.2% decrease, t(120) = 5.900, p < 0.0001, Fig. 1I, table S5), and reduced symbiont density (18.6 ± 2.5% decrease, t(48) = 7.315, p < 0.0001, Fig. 1J, table S6). In contrast to symbiotic animals, aposymbiotic Aiptasia exposed to heat, nitrate, and heat + nitrate showed no significant difference from controls across morphological, physiological, and immune metrics, with the exception of a minor reduction in total protein content under nitrate alone (12.5 ± 3.1% decrease, t(48) = 3.264, p = 0.0106, Fig. 1B–J, table S1).

### Aiptasia displayed increased canonical stress responses during heat and nitrate stress, with greater transcriptional plasticity in symbiotic than aposymbiotic animals

To identify the molecular basis of heat- and nitrate-induced physiological changes, we profiled host gene expression in both symbiotic and aposymbiotic Aiptasia across all four treatments and three genetic backgrounds. Heat exposure, with or without nitrate, induced significant shifts in host transcriptional profiles in animals of both symbiotic states (p < 0.001, Fig. 2A). Notably, symbiotic Aiptasia showed significantly greater gene expression plasticity than their aposymbiotic counterparts under heat, regardless of nitrate enrichment (Fig. 2B). Both aposymbiotic and symbiotic Aiptasia upregulated several known stress-response genes under heat stress relative to controls, including genes encoding heat shock proteins (e.g., *HSP7C* and *HSP97*), antioxidant enzymes (e.g., *PXDNL*/Peroxidasin Like), and detoxification enzymes (e.g., *GPX7*/glutathione peroxidase and *SODC.2*/superoxide dismutase) (FDR < 0.05 for all, Fig. 2C). These genes were also upregulated in symbiotic Aiptasia exposed to heat + nitrate compared to controls, along with several additional antioxidant (e.g., *PXDNL*, *PXDN.1*, *PXDNL.1*) and detoxification (e.g., *GST*/glutathione S-transferase, *GPX7*, *SODC.3*) genes (Fig. 2C). Aposymbiotic Aiptasia responded similarly, with heat and heat + nitrate leading to the upregulation of *HSP7C*, *HSP97*, *HSP7C.1*, *PXDN.1*, *PXDNL.1*, *GST* and *GPX7* (Fig. 2C). Notably, *ROMO1* (Reactive Oxygen Species Modulator 1)—a regulator of reactive oxygen species production—was consistently upregulated in all heat treatments in both symbiotic states, and symbiotic Aiptasia expressed this gene at levels 50–70% higher than observed in aposymbiotic Aiptasia, with particularly high expression under heat + nitrate (Fig. 2C).

**Fig 2.**
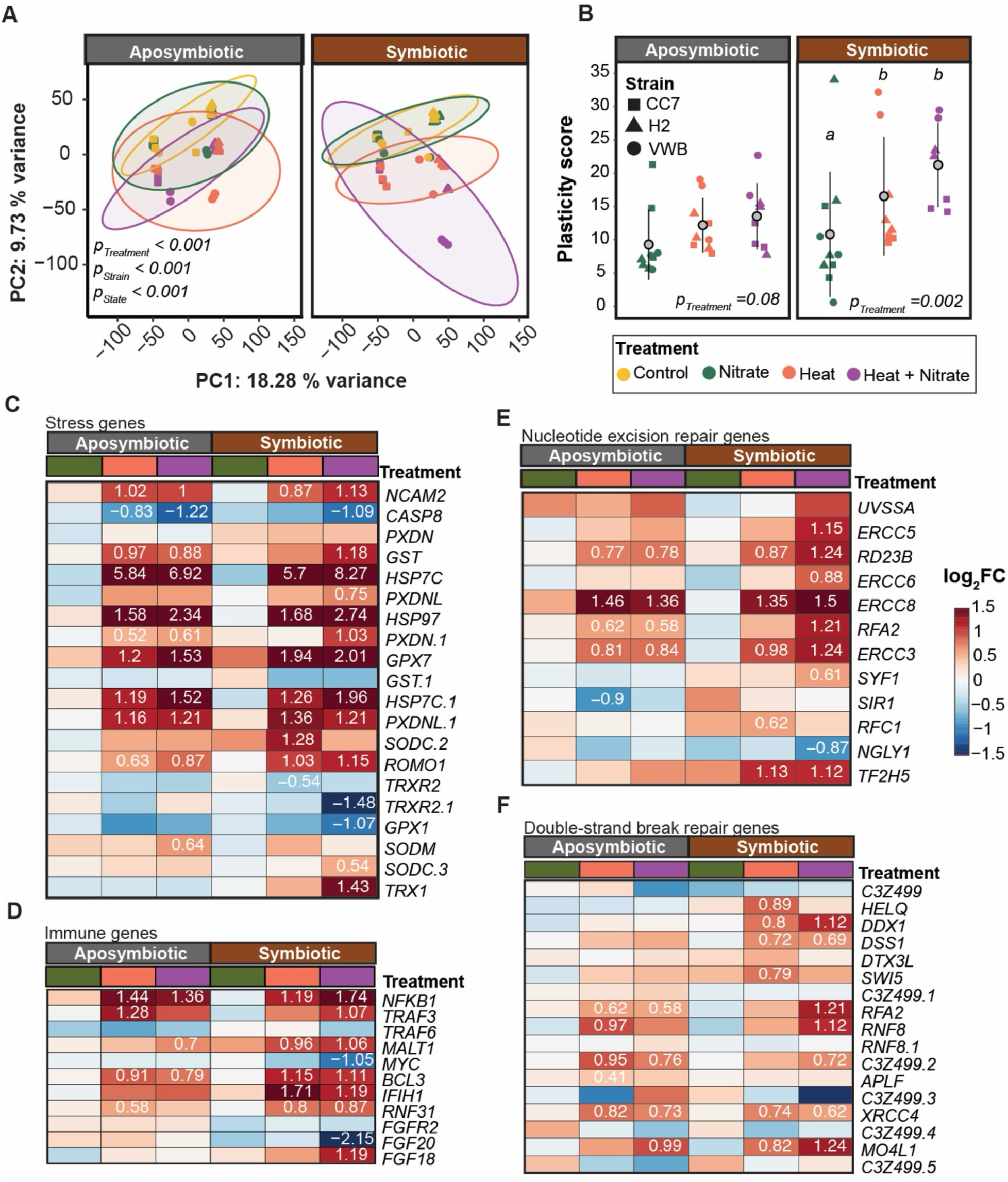
Expression of stress, immune, nucleotide excision repair, and double-strand break genes across treatments and symbiotic states. (A) Principal component analysis (PCA) of log_2_ transformed read counts for individual anemones (points) between symbiotic states. Axes indicate the percentages of variance explained by PC1 and PC2. Inset text indicates significant p-values for treatment, strain, and symbiotic state determined via a permutational analysis of variance. (B) Gene expression plasticity score for individual anemones (points) across treatments calculated using the PC distance across treatments relative to control conditions. Grey circles show the mean and black bars indicate standard deviation. Colors represent treatments (yellow = control, green = nitrate, orange = heat, purple = heat + nitrate); panel header colors denote symbiotic state (grey = Aposymbiotic, brown = Symbiotic); shapes indicate strains (square = CC7, triangle = H2, circle = VWB). Distinct letters indicate significant differences in means between treatments as determined via Tukey’s HSD post hoc tests. (C–F) Heatmaps of genes from the weighted gene co-expression network analysis brown module (fig. S2), involved in (C) stress, (D) immunity, (E) nucleotide excision repair, and (F) double-strand break repair. Rows represent genes and cell colors indicate log_2_(fold change) relative to control conditions (blue = downregulated, red = upregulated). Text in boxes denotes statistically significant (FDR < 0.05) log_2_(fold change) values. Colored boxes above heatmaps represent treatments (green = nitrate, orange = heat, pink = heat + nitrate) or symbiotic state (grey = aposymbiotic, brown = symbiotic).

To determine whether and how treatment and symbiotic state impacted immunity, we next investigated the expression of innate immunity genes. In both aposymbiotic and symbiotic Aiptasia, heat treatment led to increased expression of *NFKB1*, *MALT1* (mucosa-associated lymphoid tissue lymphoma translocation gene 1), and *BCL3* (B-cell lymphoma 3) in response to heat (FDR < 0.05, Fig. 2D). These same genes, as well as *TRAF3* (TNF Receptor Associated Factor 3), showed increased expression in symbiotic animals under heat + nitrate. A pattern-recognition receptor, *IFIH1* (Interferon Induced with Helicase C Domain 1), was also increased in symbiotic Aiptasia under both heat and heat + nitrate treatments, but was unchanged in aposymbiotic Aiptasia (FDR < 0.05, Fig. 2D).

Examination of nucleotide-excision repair (NER) genes revealed that, under heat alone, symbiotic Aiptasia upregulated *ERCC8*, *ERCC3*, and NER factors (e.g., *RFC1*) (FDR <0.05, Fig. 2E). Under heat + nitrate in symbiotic Aiptasia, a broader NER response was observed, including upregulation of several additional genes such as *ERCC5*, *ERCC6*, *ERCC8*, *ERCC3*, *RFA2*, and *SYF1* (FDR < 0.05, Fig. 2E). NER response was also observed in aposymbiotic Aiptasia, which displayed upregulation of fewer NER genes than their symbiotic counterparts under heat and heat + nitrate (e.g., *ERCC8*, *RFA2*, *ERCC3*, *RD23B*, FDR <0.05, Fig. 2E). Symbiotic Aiptasia also displayed increased expression of double-strand break (DSB) repair genes under heat relative to controls, including *HELQ*, *DDX1*, and *XRCC4* (FDR < 0.05, Fig. 2F). Under heat + nitrate, symbiotic Aiptasia displayed increased expression of additional DSB repair genes compared to controls including *RFA2*, *RNF8*, *DDX*, *MO4L1*, and *C3Z499.2* (FDR < 0.05, Fig. 2F). Aposymbiotic Aiptasia also showed increased DSB-repair gene expression under both heat and heat + nitrate treatment compared to controls (e.g., *RFA2*, *RNF8*, *C3Z499.2*, *XRCC4*, FDR < 0.05, Fig. 2F).

### Heat, but not nitrate alone, leads to ER stress and disrupted proteostasis

Transcriptomes of heat-treated symbiotic and aposymbiotic Aiptasia revealed activation of ER stress and the unfolded protein response (UPR) (Fig. 3A). For example, symbiotic Aiptasia under heat displayed increased expression of several UPR marker genes such as *GRP78* (Glucose-Regulated Protein 78), *IRE1A* (Inositol-Requiring Enzyme 1 Alpha), *PDIA4,* and *PDIA6* (Protein Disulfide Isomerase Family A Member 4 and 6) (FDR < 0.05 for all, Fig. 3A). The levels of these genes were also increased in symbiotic anemones exposed to heat + nitrate, in addition to other ER-stress modulators including *ERN2* (Endoplasmic Reticulum To Nucleus Signaling 2) and *DERL1* (Derlin-1) (FDR < 0.05, Fig. 3A). Aposymbiotic Aiptasia also displayed increased levels of ER stress genes (e.g., *GRP78*, *PDIA4*, and *PDIA6*) (FDR < 0.05, Fig. 3A) under heat and heat + nitrate, however, fewer of these genes showed increased expression in these animals as compared to symbiotic Aiptasia.

**Fig 3.**
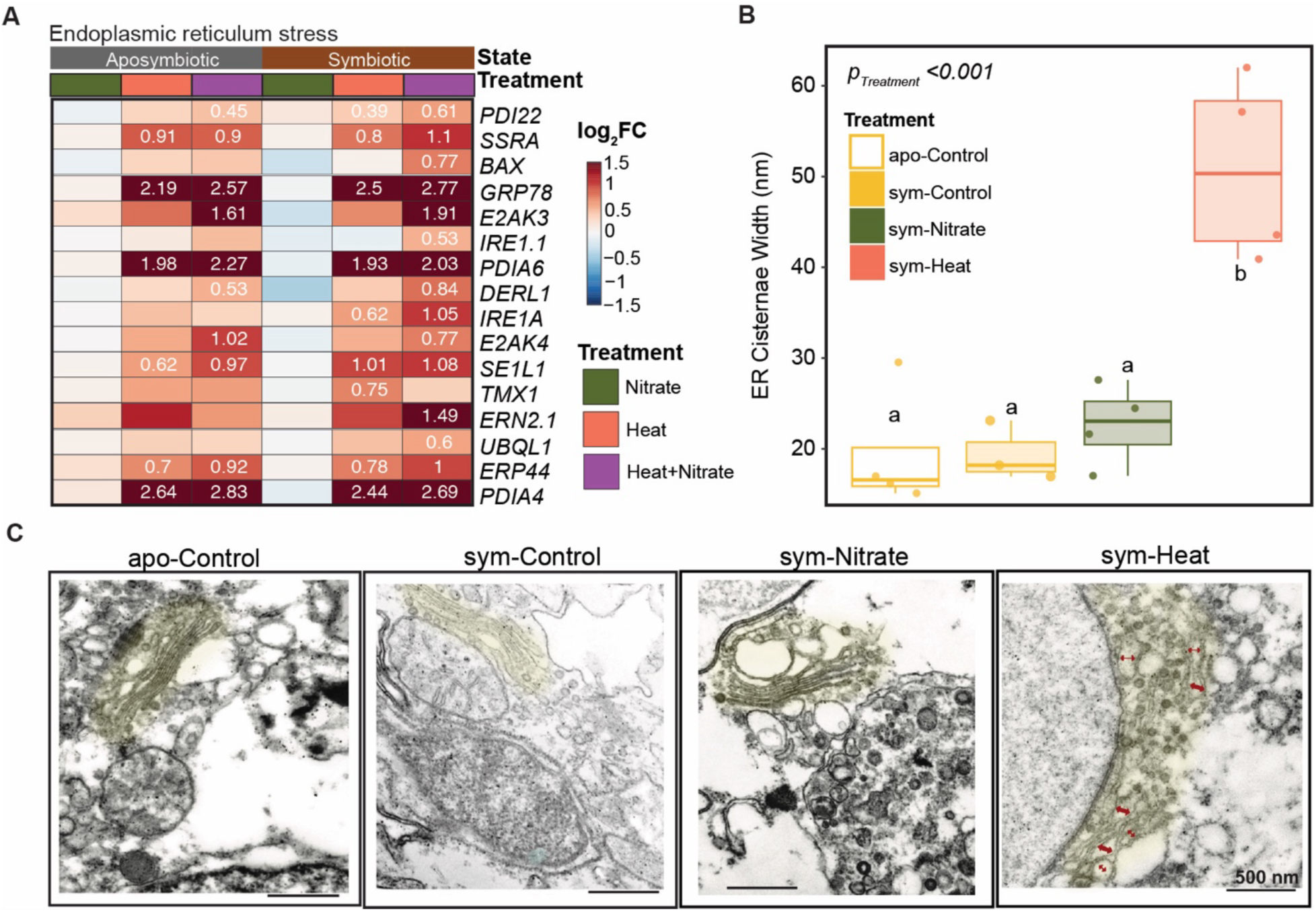
Endoplasmic reticulum (ER) stress across treatments and symbiotic states. (A) Heatmap of genes involved in endoplasmic reticulum stress from the weighted gene co-expression network analysis (fig. S2). Rows in the heatmap represent genes and the cell colors indicate log_2_(fold change) relative to control conditions (blue = downregulated, red = upregulated). Text in boxes denotes statistically significant (FDR < 0.05) log_2_(fold change) values. Colored boxes above heatmaps represent treatments (green = nitrate, orange = heat, purple = heat + nitrate) or symbiotic state (grey = aposymbiotic, brown = symbiotic). (B) ER cisternae width and (C) representative TEM micrographs of ER dilation in aposymbiotic and symbiotic Aiptasia from control, nitrate, and heat treatments. In each micrograph, the ER is highlighted in yellow, and red arrows indicate ER dilation (scale bar indicates 500 nm).

As heat was the consistent driver of proteostatic and ER stress gene expression across treatments, heat-treated animals were subjected to TEM to further investigate this response by characterizing ER morphology. Symbiotic Aiptasia under heat exhibited significantly increased ER dilation (i.e., cisternal gap widths > 50 nm) relative to controls (t(11) = −5.91, p < 0.001, Fig. 3B–C). This ER mophological response was not observed in symbiotic Aiptasia exposed to nitrate alone (t(11) = −0.61, p = 0.93, Fig. 3B–C).

### Tissue-specific expression of NF-κB protein differs between symbiotic states and under heat with or without nitrate

Due to the increases of *NFKB1* mRNA under heat stress (Fig. 2D), we next used Western blotting of whole anemone tissue lysates to determine whether protein levels of Aiptasia NF-κB (Ap-NF-κB) showed similar changes. As we have shown previously^7^, Ap-NF-κB protein levels were lower in symbiotic Aiptasia controls relative to aposymbiotic controls. The low levels of Ap-NF-κB in symbiotic anemones increased ∼25-fold under heat and heat + nitrate (F(2,9) = 18.1, p < 0.001; control vs heat: t(9) = −3.74, p = 0.012; control vs heat + nitrate: t(9) = −5.95, p < 0.001). In aposymbiotic Aiptasia, both heat and heat + nitrate treatments increased Ap-NF-κB levels ∼1.6-fold relative to controls (F(2,9) = 16.9, p < 0.001; control vs heat: t(9) = −4.94, p = 0.002; control vs heat + nitrate: t(9) = −5.13, p = 0.002, Fig. 4A–B). Although the fold change was less in aposymbiotic than symbiotic anemones, the total amount of Ap-NF-κB was similar under heat and heat + nitrate in both symbiotic states since the basal levels of Ap-NF-κB were much higher in aposymbiotic vs. symbiotic animals.

**Fig 4.**
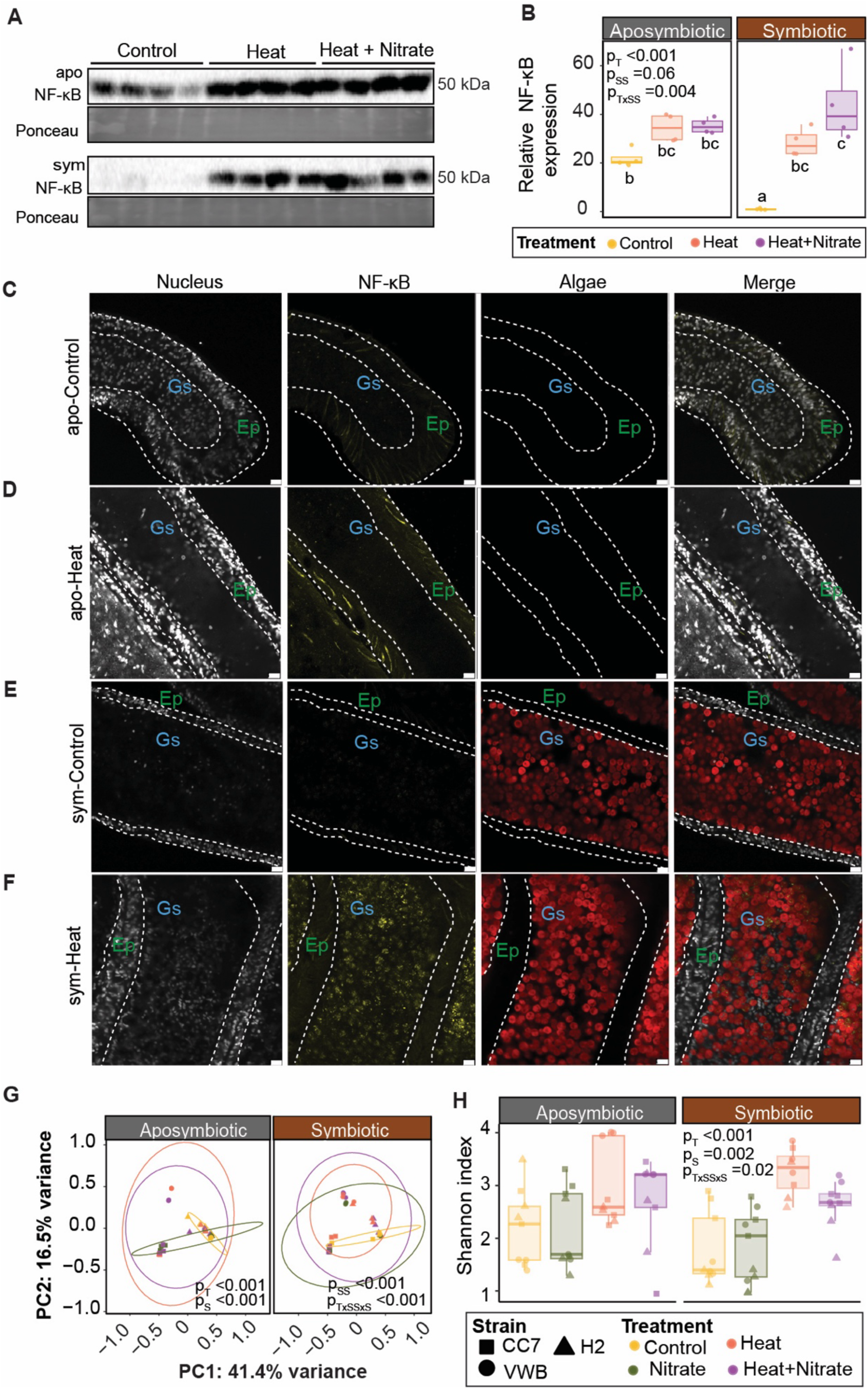
Ap-NF-κB protein expression and immunolocalization across symbiotic states and treatments. (A) Western blots of processed Ap-NF-κB (∼50 kDa) in aposymbiotic (apo) and symbiotic (sym) Aiptasia from control, heat, and heat + nitrate (H + N) treatments, with corresponding Ponceau total protein stain. Each lane contained protein from a single animal. (B) Western blot band intensity of Ap-NF-κB normalized to Ponceau staining derived from (A). The levels of Ap-NF-κB are relative to the average for control animals for each symbiotic state. Relative Ap-NF-κB levels were calculated as the ratio of Ap-NF-κB to Ponceau S signal for each sample and were then normalized to the symbiotic control average (set to 1.0). Inset text indicates p-values for treatment (T) and symbiotic state (SS) from a corresponding analysis of variance, and brackets with asterisks indicate statistically significant (p < 0.05) pairwise differences in means. (C–F) Representative confocal images of anemone tentacle tissue stained for Ap-NF-κB (yellow) and nuclei (grey), with red representing autofluorescence of symbiont chlorophyll. In all confocal images, white dotted lines denote divisions between epidermal (Ep) and gastrodermal (Gs) tissue layers, and scale bars represent 10 µm. (H) Principal coordinates analysis (PCoA) of microbiome communities based on Bray-Curtis dissimilarity, with inset text indicating significant p-values for treatment (T), strain (S), symbiotic state (SS), and interactions of these factors as determined via a permutational analysis of variance. (I) Shannon index (alpha-diversity) of bacterial communities for individual anemones (points) by strain (shape), treatment (color), and symbiotic state (panel). Inset text indicates statistical significance of linear model terms as determined via an analysis of variance.

Immunofluorescence localization of Ap-NF-κB in anemone tissues showed that the observed increases in protein levels were tissue-specific in animals of different symbiotic states (Fig. 4C–G, fig. S3–7). Specifically, aposymbiotic Aiptasia in the control group and those exposed to heat showed expression of Ap-NF-κB primarily in the epidermal layer (Fig. 4C–D). By contrast, Ap-NF-κB was visibly absent from tissues of symbiotic animals under control conditions, but was readily visible in the gastrodermal tissue layer of these animals following exposure to heat (Fig. 4E–F, fig. S7).

### Symbiosis does not influence stress-induced microbiome restructuring across treatments

To understand how heat and nitrate shaped Aiptasia bacterial microbiomes, we performed sequencing of 16S rRNA genes amplified from whole animal tissues. Principal coordinates analysis (PCoA) showed that heat treatment, with or without nitrate, significantly altered microbiome composition in both aposymbiotic and symbiotic Aiptasia (ADONIS p < 0.001, Fig. 4H). Microbial alpha-diversity (Shannon index) also increased significantly under heat and heat + nitrate for animals of both symbiotic states and strains (p_Treatment_ < 0.001, p_Strain_ = 0.002, p_T×St×SS_ = 0.002, Fig. 4I). These differences were primarily driven by heat, with both aposymbiotic and symbiotic Aiptasia showing decreased abundances of Pseudomonadales and increased abundances of Caulobacterales, Enterobacterales, Rhodobacterales, and Rhodospirillales under both heat and heat + nitrate (fig. S8). By contrast, nitrate alone had minimal effects on bacterial community composition (fig. S8).

### Symbiosis promotes cell-cycle arrest during heat stress

We next examined the expression of key cell-cycle genes, with particular interest in cell cycle progression and arrest, across treatments and symbiotic states (Fig. 5A). In symbiotic Aiptasia, heat treatment led to upregulation of *Mcm3* (Minichromosome maintenance protein 3), and downregulation of *Cdc14A_phos* (Cell Division Cycle 14A), a phosphatase involved in mitotic exit (FDR < 0.05, Fig. 5A). Similarly, symbiotic Aiptasia exposed to heat + nitrate displayed upregulation of *Mcm3*, downregulation of *Cdc14A_phos*, and upregulation of other genes such as the E2F cofactor *Dp-1-2*, *Mob1* (Mps One Binder), and the DNA-damage sensor *GADD45* (Growth Arrest and DNA Damage-inducible 45) (FDR < 0.05, Fig. 5A). In aposymbiotic Aiptasia, genes involved in activation of proliferation and DNA-damage surveillance such as *Dp-1-2*, *Mcm3*, *GADD45*, and *Mdm2* (p53-regulating ubiquitin ligase) were upregulated under both heat and heat + nitrate (FDR < 0.05, Fig. 5A).

**Fig 5.**
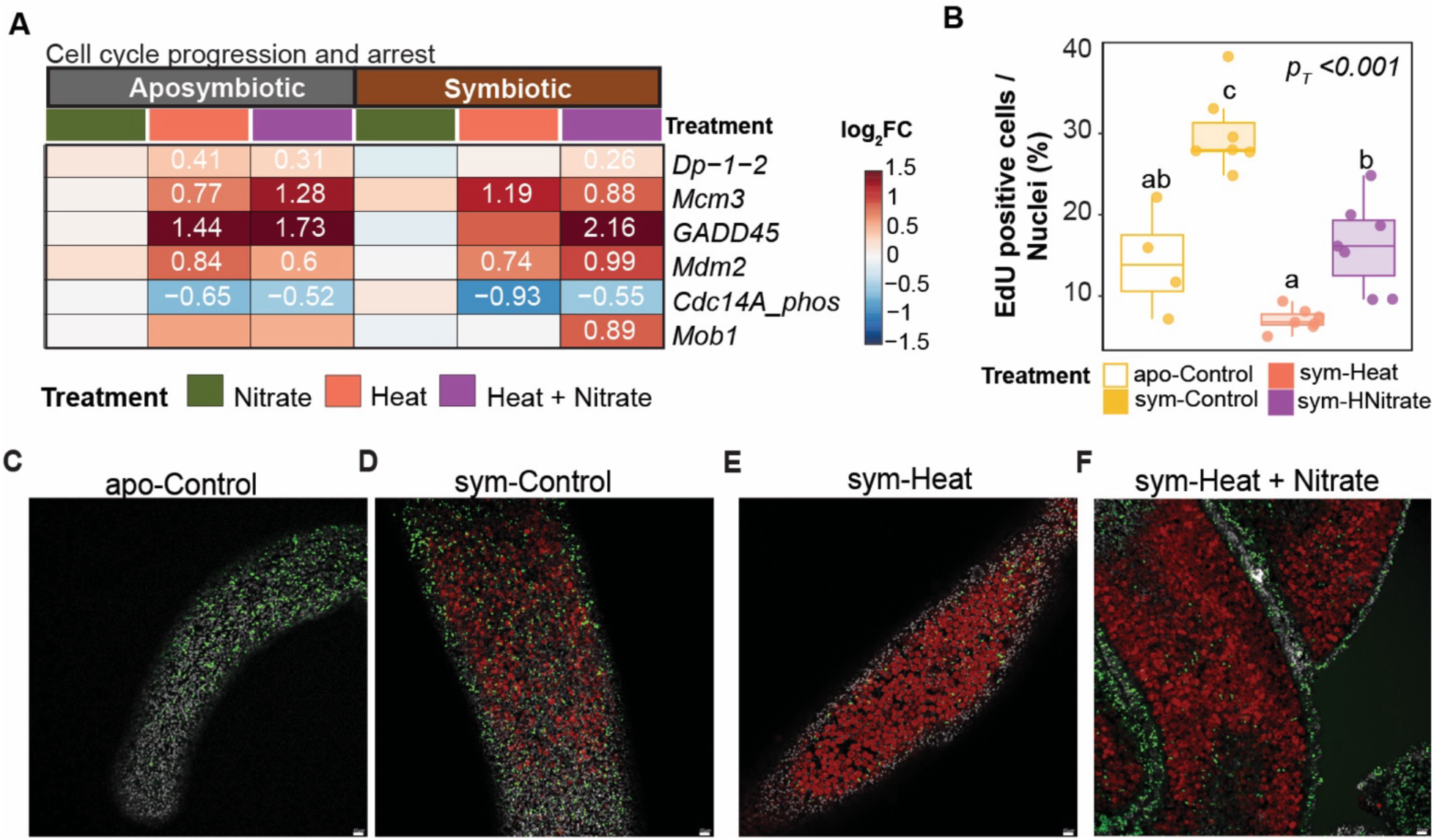
Cell-cycle gene expression and cell proliferation labeling via EdU staining across symbiotic states and treatments. (A) Heatmap of genes involved in cell cycle progression and arrest. Rows represent genes and cell colors indicate log_2_(fold change) relative to control conditions (blue = downregulated, red = upregulated). Text in boxes denotes statistically significant (FDR < 0.05) log_2_(fold change) values. Colored boxes above heatmaps represent treatments (green = nitrate, orange = heat, purple = heat + nitrate) or symbiotic state (grey = aposymbiotic, brown = symbiotic). (B) Quantification of EdU positive cells relative to nuclei positive cells by treatment. Inset text indicates p-values for treatment (T) from a corresponding analysis of variance, and letters indicate statistically significant (p < 0.05) pairwise differences in means. (C–F) Representative confocal images of anemone tentacle tissue stained for EdU (green) and nuclei (grey), with red representing autofluorescence of symbiont chlorophyll. Scale bars represent 10 µm.

To determine whether the transcriptional reprogramming of cell-cycle genes had functional consequences for host cell proliferation, we used EdU incorporation assays in symbiotic Aiptasia (Fig. 5B–F). Since aposymbiotic anemones showed less apparent gross morphological and physiological responses to heat, we focused on symbiotic animals in which disruption of cell-cycle gene expression was most pronounced. Heat exposure significantly reduced EdU labeling (76.4 ± 8.3%, t(21) = 9.23, p < 0.001), and while heat + nitrate also reduced proliferation, the effect was less severe (45.7 ± 8.3%, t(21) = 5.53, p = 0.0001, Fig. 5B–F).

### Heat and nitrate reduced host-symbiont nutrient exchange

To assess the impact of heat, nitrate, and their combination on host-symbiont nutrient exchange, we first assessed the expression of genes previously identified as candidates involved in this process based on expression in aposymbiotic vs. symbiotic Aiptasia^16^ (FDR < 0.05, Fig. 6B). Both aposymbiotic and symbiotic Aiptasia showed decreased levels of genes involved in sterol transport and nitrogen shuttling under heat and heat + nitrate, including *NPC1.1* (Niemann-Pick Disease Type C1) and *AMT1.1* (Ammonium Transporter 1) (FDR < 0.05, Fig. 6B). Moreover, heat + nitrate exposure was associated with upregulation of *G3P* (Glyceraldehyde 3-phosphate) and downregulation of *GPDM* (mitochondrial Glycerol-3-Phosphate Dehydrogenase) relative to heat alone in symbiotic anemones (FDR < 0.05, Fig. 6B). In aposymbiotic animals, *G3P* was increased only in heat-treated anemones, whereas *RPE* (Ribulose-5-Phosphate-3-Epimerase) was increased under heat + nitrate, whereas *GPDM* was decreased under both treatments (FDR < 0.05, Fig. 6B).

**Fig 6.**
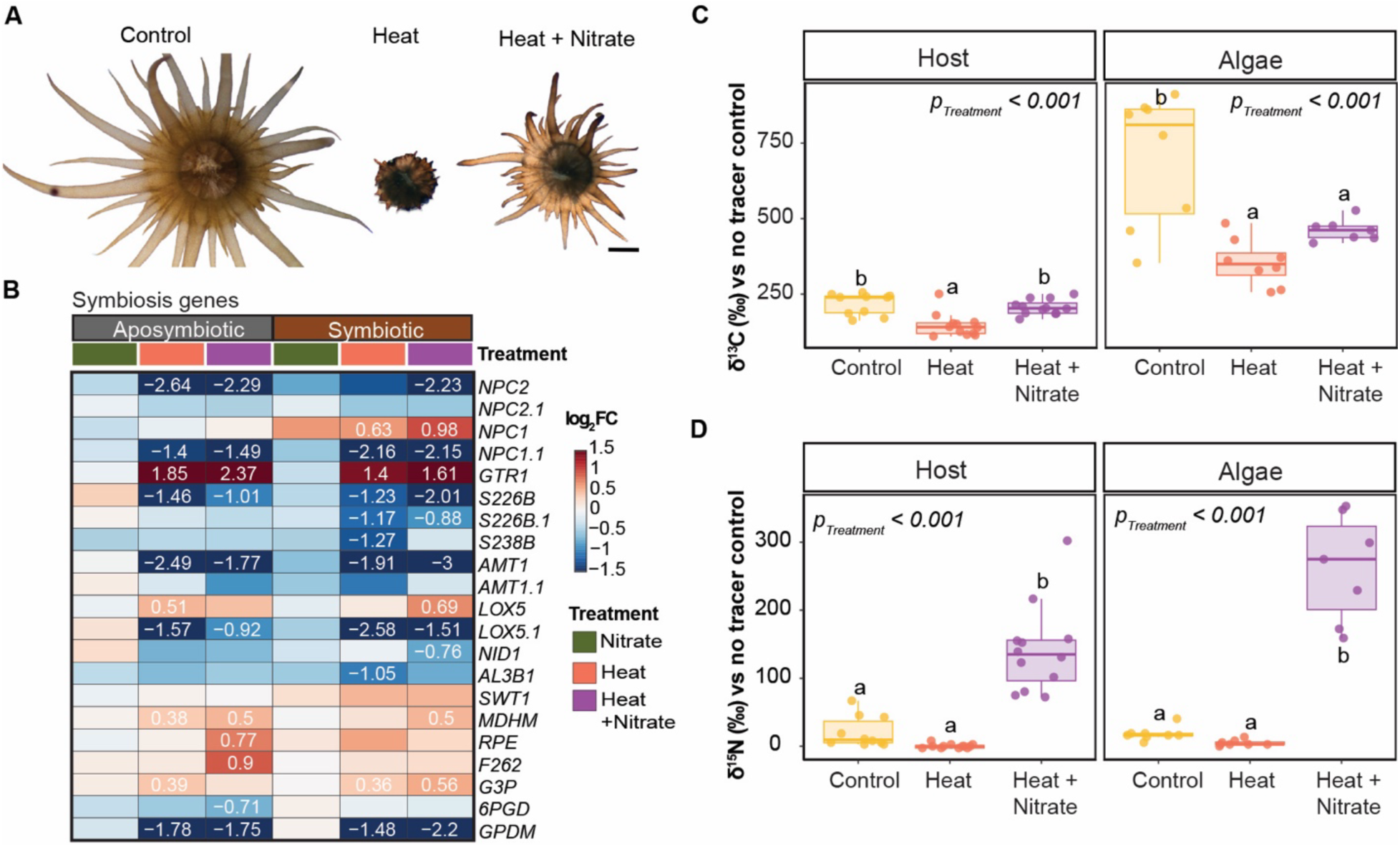
Regulation of host-algal symbiont nutrient dynamics across treatments. (A) Representative photos of symbiotic Aiptasia across treatment at experimental day 12. (B) Heatmap of differentially expressed genes involved in host-algal symbiont nutrient exchange. Rows represent genes and cell colors indicate log_2_(fold change) relative to control conditions (blue = downregulated, red = upregulated). Text in boxes denotes statistically significant (FDR < 0.05) log_2_(fold change) values. Colored boxes above heatmaps represent treatments (green = nitrate, orange = heat, purple = heat + nitrate) or symbiotic state (grey = aposymbiotic, brown = symbiotic). (C–D) Bulk isotope measurements of (C) δ13C and (d) δ15N for host and algal symbiont fractions from individual symbiotic anemones (points) across treatments, normalized to sample fractions without tracers. Black bars indicate standard deviation. Colors represent treatments (yellow = control, orange = heat, purple = heat + nitrate). Inset text indicates p-values for treatment (T) from a corresponding analysis of variance, and letters indicate statistically significant (p < 0.05) pairwise differences in means.

To further explore how heat and nitrate impact nutrient dynamics, we measured bulk carbon and nitrogen isotope composition (δ^13^C and δ^15^N) in host and algal symbiont fractions from individual symbiotic Aiptasia across treatments (Fig. 6C–D). In the host fraction, heat significantly reduced δ^13^C by 32.6 ± 6.6% relative to the control (t(31) = 4.97, p = 0.0001, Fig. 6C), whereas heat + nitrate did not significantly affect host δ13C (5.0 ± 6.6% lower than control, t(31) = 0.80, p = 0.7042). In contrast, host δ^15^N did not differ significantly under heat alone (t(31) = 1.18, p = 0.4742), but was strongly enriched under heat + nitrate (t(31) = −6.93, p < 0.0001, Fig. 6D). In the algal symbiont fraction, heat also significantly reduced δ^13^C by 49.5 ± 9.8% relative to the control, and algal symbiont δ13C remained significantly reduced under heat + nitrate (34.1 ± 10.2% lower than control, t(20) = 5.03, p = 0.0002; t(20) = 3.36, p = 0.0085, respectively, Fig. 6C). Similar to the host fraction, algal symbiont δ^15^N did not differ significantly from the control under heat alone (t(20) = 0.62, p = 0.8106), but was strongly enriched under heat + nitrate (t(20) = −10.9, p < 0.0001, Fig. 6D).

### Immune activation under heat + nitrate did not confer protection against pathogen challenge

To test whether the immune activation observed under heat + nitrate conferred improved tolerance to pathogen challenge, we focused on symbiotic anemones, which exhibited the greatest fold-increase in Ap-NF-κB levels following heat + nitrate treatment (44-fold vs. 1.6-fold in aposymbiotic animals). We exposed symbiotic anemones to a marine bacterial pathogen, *Pseudomonas aurigenosa* (Pa14), and then compared survival outcomes across treatments (Fig. 7A). Survival differed significantly among treatments (χ^2^ = 56.0, df = 4, p<0.001, Fig. 7B). By day 13, survival probability was significantly reduced in all treatment groups compared to non-pathogen-treated controls (p < 0.0001). Control anemones (i.e., not exposed to the pathogen) showed 100% survival probability, whereas heat stress alone reduced survival probability to 16.7%. The combined heat + nitrate treatment resulted in 66.7% survival, and pathogen exposure (infected) alone decreased survival to 41.7%. Notably, anemones subjected to combined heat + nitrate + pathogen exhibited complete mortality, with survival probability reaching 0% by day 8 (Fig. 7B). Taken together, these results indicate that even though heat + nitrate treatment can robustly induce NF-κB expression in symbiotic Aiptasia, this does not lead to enhanced survival in the presence of a challenge with a bacterial pathogen.

**Fig 7.**
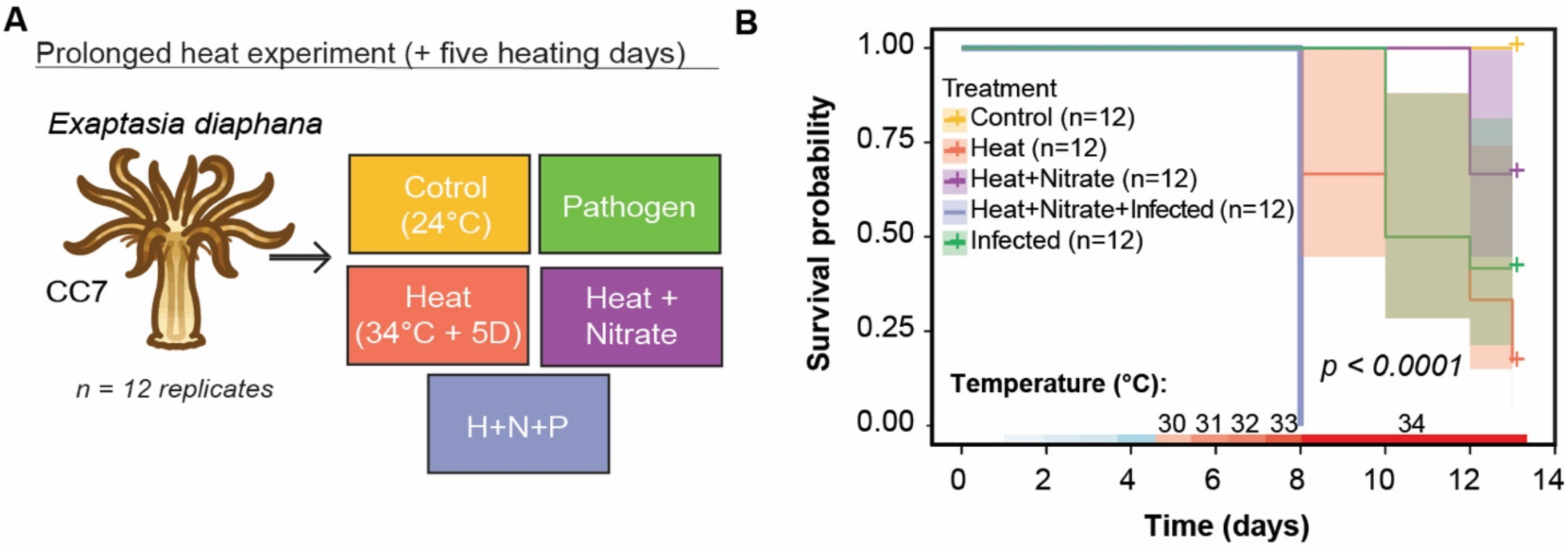
Design and survival outcomes for pathogen challenge experiments. (A) Schematic diagram of the experiment with extended heating days (ramp 1° C per day for 8 days to 34° C, then held for seven days). The marine pathogen, *Pseudomonas aeruginosa* (Pa14), was introduced when temperature reached 34°C. Colored boxes indicate treatments: yellow (control), green (pathogen), orange (heat), purple (heat + nitrate), and blue (heat + nitrate + pathogen). (B) Survival curves of symbiotic Aiptasia across treatments. No mortality was observed in control conditions. Inset text indicates p-value for treatment.

## DISCUSSION

Here, we investigated how heat and nitrate impact the Aiptasia-dinoflagellate symbiosis across biological scales, encompassing host and symbiont physiology, metabolism, gene expression, and bacterial microbiomes. These data reveal how these two stressors interact to disrupt the regulatory processes that maintain symbiosis, including host immune suppression^7^, cell-cycle control^8^, and nutrient exchange^9^. Each of these processes was disrupted by heat stress, with more pronounced responses in symbiotic than aposymbiotic Aiptasia. Notably, increased pathogen sensitivity under heat suggests that immune activation reflected organismal stress rather than improved pathogen resistance. Nitrate supplementation temporarily buffered these effects, but failed to sustain stable symbiosis under prolonged heat exposure. Together, these findings support a mechanistic model in which symbiont dysfunction and host physiological decline converge to drive redox imbalance, which acts as a key mediator of heat-induced symbiosis breakdown by destabilizing the coupled regulatory processes required to maintain symbiosis.

### Algal symbionts drive heightened host thermosensitivity

In this study, the presence of algal symbionts fundamentally altered host responses to stress, with more pronounced heat-induced physiological declines observed in symbiotic compared to aposymbiotic Aiptasia. Observations spanning morphology, physiology, and gene expression suggested a disproportionate energetic and regulatory burden for symbiotic hosts under stress compared to aposymbiotic hosts. This finding differs from what has been found in the temperate coral *Astrangia poculata*, where symbiosis was shown to reduce host stress responses under heat^32^. Differences in heat exposure regimes may explain these contrasting findings^33^, as our heat treatment was sufficiently intense to compromise symbiont performance and induce breakdown of symbiosis. This finding highlights a context-dependent trade-off inherent to metabolic integration in cnidarian-Symbiodiniaceae symbiosis, in which host-symbiont metabolic coupling becomes a burden under stress, amplifying the host stress response rather than buffering it^34^.

Under heat, the presence of symbionts elicited a pronounced oxidative stress response, linking early photophysiological impairment of symbionts to downstream declines in host health. This oxidative stress signature is typical for cnidarian holobionts under stress, with corals exposed to heat consistently showing upregulation of molecular chaperones (e.g., Heat Shock Proteins)^35,36^, antioxidant enzymes (e.g., peroxidases, glutathione s-transferases, superoxide dismutases)^37^, and proteostasis pathways (e.g., UPR/ER stress responses)^6,22,38,39^. In this study, upregulation of *ROMO1* and several antioxidant enzymes under heat in symbiotic Aiptasia was consistent with excess ROS accumulation^40^, reflecting a common feature of bleaching in which symbionts generate ROS that damage host tissues^41^. Beyond direct cellular damage, increased ROS (derived both from the host and symbionts) is expected to disrupt redox-sensitive pathways^42^, likely amplifying physiological impairment of both partners. Although we did not quantified host-vs. symbiont-derived ROS, the convergence of symbiont dysfunction, host physiological decline, and oxidative stress supports the hypothesis that redox imbalance is a key mediator of heat-driven physiological declines in symbiotic Aiptasia.

Upregulation of UPR pathway genes in both aposymbiotic and symbiotic Aiptasia suggested that heat stress imposed a proteostatic burden on hosts regardless of symbiotic state. Heat exposure can induce proteostatic stress through both direct and indirect mechanisms. For example, heat physically disrupts protein structure^43^, resulting in unfolding that inactivates protein function^43^ and promotes toxic aggregate formation^44^. At the same time, mounting a heat stress response requires extensive cellular resources (e.g., ATP)^45^, thus limiting the ability of the organism to devote resources to clear and replace unfolded proteins. Consistent with this hypothesis, heat exposure resulted in upregulation of *IRE1* and the ER chaperone *GRP78*/*BiP*, together with upregulation of three protein disulfide isomerase (*PDI*) genes, which are folding catalysts that facilitate correct disulfide bond formation during protein refolding^46^. These markers point to activation of an ER proteostasis mechanism that expands folding capacity and supports disposal of misfolded proteins via canonical UPR signaling^47^. Consistent with this hypothesis, in symbiotic Aiptasia, increased ER dilation indicated proteostatic burden^39^. Similar ER dilation in response to stress has been reported in other systems^48^ and aligns with overburdened UPR signaling^49^. These findings suggest that heat drives proteostasis disruption in Aiptasia, contributing to runaway dysbiosis and downstream host declines.

Heat also elicited a localized immune activation in symbiotic Aiptasia that reflects symbiosis destabilization rather than a coordinated host defense. In symbiotic animals, NF-κB protein levels were elevated under heat within the gastrodermal cells where algal symbionts reside. This tissue-specific analysis extended previous observations that heat-induced increases in NF-κB expression are associated with symbiont loss^7,16^. Specifically, these findings demonstrated that heat-induced increases in NF-κB protein are not whole-organism responses, but instead are localized to symbiont-containing cells where dysbiosis initiates. Our data therefore support a model in which heat drives a localized shift away from immune suppression in gastrodermal cells towards immune activation (likely in part via NF-κB), which contributes to algal loss. This interpretation aligns with prior work showing that thermal stress engages immune effector pathways that overlap with canonical defense responses. For example, the immune and defense pathway activation in thermal stress is similar to that also engaged during infection^50^, and both heat stress and pathogen exposure have been shown to induce oxidative-defense enzymes and activate the melanization pathway^50,51^, mirroring immune enzyme increases observed here under heat. Moreover, these defense-like responses — such as activation of NF-κB and other immune-associated genes — have been proposed to reflect an early damage-surveillance response that can precede symbiosis collapse^7,23,52^. Notably, elevated immune responses during pathogen infection are associated with decreased symbiont density in corals^53^, which also display induction of innate immune transcriptional programs (including NF-κB signaling) and immune enzymes during thermal stress^54^. While these findings do not prove causation, they consistently correlate increased immune activity with reduced symbiont density. Further supporting this interpretation, although NF-κB levels increased under heat + nitrate, symbiotic Aiptasia showed rapid mortality within 24 h following pathogen exposure, demonstrating that NF-κB activation did not confer functional resistance to pathogen infection^23^. This dissociation between NF-κB activation and immune competence suggests that, under heat stress, NF-κB signaling reflects a response to cellular damage and oxidative stress^16^, rather than a coordinated immune defense. This result supports our broader interpretation that heat- and nitrate-associated immune activation reflects destabilization of symbiosis, not preserved or restored host immune function.

In our experiments, heat exposure increased bacterial community alpha-diversity and reshaped composition in Aiptasia regardless of symbiotic state, suggesting that thermal stress broadly weakens host regulation of microbial communities. Our data indicate that microbiome dysbiosis under heat is likely a downstream consequence of host stress and immune dysregulation, rather than an initiating driver of holobiont collapse^55,56^. Enterobacterales and Rhodobacterales consistently increased under heat regardless of symbiotic state, mirroring previous findings that both orders of bacteria increase in corals experiencing stress and disease^57,58^. Similar shifts toward opportunistic bacterial taxa have been documented during coral bleaching events^59^, suggesting that bacterial dysbiosis is a consequence rather than a cause of holobiont destabilization^55^. These findings indicate that opportunistic shifts in microbiome diversity are a common feature of bleaching, but they do not distinguish cause from consequence.

Heat stress disrupted host cell-cycle regulation — a mechanism previously shown to regulate symbiont growth and maintain symbiosis stability^8^. This disruption induced damage-associated growth arrest, accompanied by physiological declines and a reallocation of host resources from growth to survival. In symbiotic Aiptasia, heat stress led to upregulation of minichromosome maintenance protein 3 (*Mcm3*) and downregulation of cell division cycle 14A (*Cdc14A_phos*), a phosphatase implicated in mitotic exit, consistent with disrupted cell-cycle progression^60^. Together with reduced EdU labeling, these findings indicate that heat suppresses cell proliferation, potentially impairing tissue renewal precisely when repair demands are highest. Genes implicated in DNA damage signaling and checkpoint regulation were also upregulated in symbiotic Aiptasia under heat, suggesting that high heat induced DNA damage^22^ and unresolved growth arrest^61^. Checkpoint activation occurred along with oxidative/immune stress and metabolic disruption, suggesting that growth arrest is an emergency response to damage rather than a controlled mechanism of symbiosis regulation under heat^61^. Disruption of the cell cycle likely is a downstream effect of redox/immune stress, linking molecular stress to impaired renewal and helping in part to explain why symbiotic animals experienced greater physiological costs. Overall, as symbiont function declines, symbiosis augments redox, immune, and proteostatic demands, accelerating holobiont destabilization.

### Disruption of carbon-nitrogen balance under heat contributed to symbiosis breakdown

Heat stress disrupted metabolic interactions between hosts and algal symbionts, as indicated by suppression of ammonium and sterol transport machinery and changes in carbon and nitrogen incorporation between partners. These findings are broadly consistent with previously observed heat stress responses in Aiptasia^62^ and multiple coral species^9,23^. This further supports the model proposed by Cui et al.^62^ that, under heat, the host loses its capacity to enforce nitrogen limitation (a core stabilizing feature of symbioses^63^) as carbon supply tightens, allowing nitrogen to flow toward symbiont growth rather than being retained/assimilated by the host^62^. As an effect, the algal symbiont becomes more like a parasite, using its photosynthates for its own survival, rather than continuing translocation to the host^9,24^. Further, our isotope data indicated reduced symbiont-derived carbon in the host under heat, consistent with multiple studies in corals demonstrating the ability of heat to limit host carbon^9,24^. A key implication is that heat reduces metabolic input when host energetic demand is elevated by other stress response pathways, creating a mismatch between rising energy demands and constrained energy supply. Since hosts rely heavily on symbiont-derived carbon, reduced nutrient exchange likely limits metabolic flexibility and exacerbates physiological stress.

### Heat + nitrate delayed physiological decline but did not restore regulatory control

Heat + nitrate delayed physiological decline by briefly maintaining symbiont function and host carbon acquisition, without restoring the regulatory control that supports long-term host health. In early heat + nitrate exposure, symbiotic Aiptasia exhibited delayed bleaching and transient preservation of host morphology, yet gene expression patterns remained similar to those under heat alone. These findings are consistent with prior work showing that the addition of nitrate fails to mitigate cellular stress imposed by heat^64^. Our stable isotope analyses further revealed elevated nitrogen incorporation in both symbiont and host under heat + nitrate, indicating that nitrate enrichment lifted nitrogen limitation within the holobiont. Additionally, hosts displayed continued incorporation of symbiont-derived fixed carbon under heat + nitrate, whereas heat alone was associated with carbon-limited state of the host^9^. This short-term buffering under heat + nitrate coincided with higher algal symbiont densities and improved photosynthetic efficiency compared with heat alone, indicating that the buffering effect of nitrate depends on the presence of a functional algal symbiont population^65^. However, this effect of nitrate under heat did not result in long-term host health. Despite sustained translocation of fixed carbon under heat + nitrate, host gene expression indicated that this carbon was not used for growth and tissue renewal. Instead, upregulation of glyceraldehyde-3-phosphate dehydrogenase (*G3P*) and mitochondrial malate dehydrogenase (*MDHM*) suggest that carbon facilitated the maintenance of cellular redox balance^66,67^. Downregulation of mitochondrial glycerol-3-phosphate dehydrogenase (*GPDM*) further indicated limited transfer of this carbon into mitochondrial energy production^68^, suggesting that high nutrient availability only temporarily buffers heat stress, without solving energetic deficiencies.

Our findings together support a multi-scale model as to how heat triggers dysbiosis through effects on three key biological processes. Specifically, the decline in symbiont function under heat stress amplifies the host’s stress responses, thereby disrupting crucial processes including immune regulation, cell-cycle control, and nutrient exchange. While nitrate enrichment can temporarily mitigate visible organismal decline by maintaining symbiont function and metabolic input, it fails to restore the fundamental regulatory mechanisms necessary for the long-term stability of symbiosis during prolonged heating. Overall, these findings highlight the need for intervention to minimize local and global stressors, in addition to heat, that threaten cnidarian-algal symbioses.

## MATERIALS AND METHODS

### Aiptasia culture

Anemones (*Exaiptasia diaphana*) used in these experiments were of three laboratory strains: CC7^69^, H2 and VWB^70^. Animals were maintained in long-term (> 2 yr) culture as either symbiotic (hosting homologous algal strains) or aposymbiotic, with aposymbiotic animals produced via menthol bleaching of symbiotic anemones^71^. Prior to the experiment, 48 individuals per strain per symbiotic state (N = 144 animals per symbiotic state) were transferred to individual wells of six-well plates with 10 mL of filtered seawater (FSW). These plates were placed in an incubator at 24°C (± 1°C, mean ± SD) under white fluorescent light (40 µmol photons m⁻² s⁻¹) on a 12-hour light:12-hour dark cycle (light from 10:00-22:00). Animals were fed twice weekly via the addition of 50 μL of a homogenous mixture of freshly hatched *Artemia salina* nauplii in FSW to each well. Wells were cleaned 30 min after feeding using cotton swabs, followed by a complete water change. Animals were acclimated under these conditions for 7 days prior to use in the experiment.

### Multistressor mesocosm experiment

All glassware was acid-washed with 10% v/v hydrochloric acid and thoroughly rinsed with deionized water. Acclimated anemones were assigned to one of four treatment groups: control, heat, nitrate, or heat + nitrate (Fig. 1A). Each treatment group comprised 12 anemones per strain per symbiotic state (N = 36 animals per symbiotic state per treatment). The treatments lasted for a total of 14 days (Fig. 1A). First, all animals were placed in one of two incubators at 25°C under lights as described above. The incubator containing the control and nitrate groups was maintained at 25°C throughout the experiment, while the incubator containing the heat and heat + nitrate groups was ramped at a rate of 1°C per day for 10 days to 34°C. Beginning on day 5 for the nitrate and heat + nitrate groups, animals were kept in FSW supplemented with 5 μM NH_4_NO_3_, which was prepared fresh daily. All anemones were fed freshly hatched *Artemia salina* nauplii every other day, and complete water changes were performed 2 h following feeding. At the end of the experiment (day 12), anemones were flash-frozen in liquid nitrogen and stored at −80°C for subsequent processing.

### Quantification of seawater nutrient concentrations

On days 5–9, prior to water changes, seawater was collected from the stock solution of FSW supplemented with 5 µM ammonium nitrate (NH_4_NO_3_) for nutrient analysis. Specifically, a 50 mL sample was drawn using a 60 mL acid-washed syringe, filtered through a glass fiber filter (Whatman GF/F, 0.7 µm) into an acid-washed, deionized-leached polyethylene bottle, and frozen (−18°C) prior to analysis. Later, samples were thawed and the concentration of NO_3_^-^ was determined using high-resolution digital colorimetry on a SEAL Nutrient AutoAnalyzer 3 with segmented flow injection using standard techniques^72–74^.

### Morphological and physiological assessments

Morphological observations were conducted daily at 15:00 to avoid interfering with feeding schedules. For each animal in the experiment (N = 36 per symbiotic state per treatment), tentacle phenotypes were recorded as 1 (dome shape: compressed body with closed tentacles), 2 (slightly compressed body with shortened tentacles), 3 (rounded body with short, extended tentacles), or 4 (fully-extended body with extended tentacles). Polyp extension scores were adapted from^75^. No anemone mortality was observed throughout the experiment.

Dark-adapted photochemical efficiency (F_v_/F_m_) was also measured daily in symbiotic and aposymbiotic individuals using a Walz Junior-PAM™. Measurements were taken in triplicate per individual (N = 24 animals per symbiotic state per treatment) from different sections of the oral disc. Calibration was performed using either FSW or FSW with 5 µM NH_4_NO_3_ to match culture conditions.

Anemones flash-frozen at the end of the treatments were used to quantify several physiological metrics. First, animals were homogenized with glass beads using a Tissue-Tearor (Model 985370, BioSpec Products, Inc.) at a speed of 6 m/s for 2 min. The resulting tissue slurry was centrifuged at 3,300 rpm for 3 min at 4°C to separate host (i.e., anemone) and algal symbiont fractions. Total host protein content (N = 9 animals per symbiotic state per treatment) was quantified using a Bradford protein assay^76^, with each sample run in triplicate and concentrations determined using a bovine serum albumin standard curve. Absorbance was measured at 595 nm using a microplate spectrophotometer (Biotek Synergy H1; CA, USA). Total host carbohydrates (N = 9 animals per symbiotic state per treatment) were measured using the phenol-sulfuric acid method^77^ with absorbances read at 500 nm, and concentrations (mg mL^-1^) were calculated using a D-glucose standard curve before being normalized to total host protein.

To extract host protein for immune enzyme assays, anemones were homogenized in 500 µL of cold cell lysis buffer (RIPA buffer: 100 mM Tris, 100 mM NaCl, 10 mM EDTA, and 1x protease inhibitor cocktail) using a micropestle. The resulting tissue slurry was centrifuged at 14,000 x *g* for 15 min at 4°C, then the supernatant was transferred to a new prechilled 1.5 mL microcentrifuge tube and immediately used in enzyme assays. Peroxidase (POX) activity (N = 9 animals per symbiotic state per treatment) was measured by combining 10 µL of protein extract with 40 µL of sodium phosphate buffer (10 mM, pH 6.0) and 50 µL of guaiacol (25 mM in 0.01 M phosphate buffer) in a 96-well plate, with 3 technical replicates per sample. Next, 10 µL of hydrogen peroxide (20 mM in 0.01 M phosphate buffer) was added, and the absorbance at 470 nm was measured every 30 sec for 15 min. POX activity was calculated as the change in absorbance per mg protein (ΔA490*mg^-1^*min^−1^) (Mydlarz and Harvell 2007). Phenoloxidase activity (N = 9 animals per symbiotic state per treatment) was quantified by mixing 20 μL of sodium phosphate buffer (50 mM, pH 7.0), 25 μL of sterile water, and 20 μL of the protein extract, with 2 technical replicates per sample. Dopamine (30 μL, 10 mM) was added as the substrate, and the absorbance at 490 nm was measured every 30 sec for 15 min. The change in absorbance was calculated during the linear range of the curve (∼1–3 min), and activity was expressed as the change in absorbance per mg protein (ΔA490*mg^-1^*min^-1^)

Algal symbiont densities (N = 9 animals per symbiotic state per treatment; expressed as algal cells per mg of host protein) were determined by analyzing 10 μL aliquots of algal slurry using a hemocytometer under a light microscope, with three technical replicates per sample. Replicate counts were averaged, extrapolated to the total slurry volume, and then normalized to the total host protein concentration.

### Statistical analysis and visualization of morphological and physiological data

Analyses were performed using R v4.4.0^78^, and visualizations were generated using *ggplot2*^79^. To compare physiology between aposymbiotic and symbiotic anemones across treatments, data were confirmed to meet relevant assumptions for parametric statistics, then l linear models were built relating each metric to the fixed interactive effect of treatment, strain, and symbiotic state (metric ∼ treatment*strain*state). Analyses of variance (ANOVAs) were conducted to determine significance levels of the model terms. Where appropriate, the significance levels of pairwise comparisons of group means were determined using Tukey’s Honest Significant Difference (HSD) post-hoc tests. Results of these statistical analyses and comparisons are reported in table S1–6. For visualization, relative changes in physiological traits were calculated by dividing measurements for each animal by the average of the measurements for the animals of the same symbiotic state in the control group. Further, to evaluate whether treatment influenced overall physiological profiles, principal component analyses (PCA) were performed on data separated by symbiotic state. The significance level of each term (treatment, strain, and treatment*strain) in the PCA models was determined using a permutational analysis of variance (PERMANOVA; Bray-Curtis dissimilarity matrix, 999 permutations) performed using the adonis2() function from *vegan*^80^.

### Gene expression profiling and weighted gene co-expression network analysis

Total RNA was isolated from anemones (N = 9 per treatment per symbiotic state, 72 animals total, using an RNAqueous Total RNA Isolation Kit (Invitrogen) according to the manufacturer’s protocol, with an additional DNA removal step using the DNA-free™ DNA Removal Kit (Invitrogen). RNA was quantified on a Denovix spectrophotometer and run on a 1% agarose gel to confirm RNA integrity. Normalized RNA extracts (20 ngµL^-1^ per sample) were sent to the University of Texas at Austin’s Genome Sequencing and Analysis Facility. TagSeq libraries were prepared following^81^, and single-end (100 bp) sequencing was performed on the NovaSeq6000 platform (Illumina).

Demultiplexed reads for each sample were trimmed using Fastx_toolkit. Briefly, 5’-Illumina leader sequences, poly (A)+ tails, and sequences that were less than 20 bp in length with < 90% of bases having quality cutoff scores < 20 were removed. PCR duplicates were assessed and removed from all libraries. After quality filtering, reads were mapped to the *E. diaphana* genome^82^ using *bowtie2*^83^, and a custom Perl script was used to generate a summary count table for downstream analysis. All read tracking statistics are provided in table S7. Only host data were analyzed, as not enough symbiont reads were recovered for robust analysis.

Differentially expressed genes (DEGs) were identified using *DESeq2*^84^ using the model: design = ∼ treatment + strain + symbiotic state. Genes were considered differentially expressed if the corresponding treatment contrasts (e.g., heat vs. control) had an FDR-adjusted p-value < 0.05. Counts were rlog-transformed in DESeq2 and visualized via a PCA using the prcomp() function. The effect of treatment on gene expression profiles was tested using a PERMANOVA as described above. Overall gene expression plasticity between the heated treatment relative to control conditions of each symbiotic state was determined by calculating the weighted distance between samples along the first two PC axes following^85^. Differences in plasticity were assessed using a linear model and ANOVA with Tukey’s HSD post-hoc tests as described above.

A weighted gene co-expression network analysis (WGCNA) was performed to identify modules of co-regulated genes. A signed network was constructed using a soft threshold power of 10, a minimum module size of 30, and a module merging threshold of 40% dissimilarity. Module-trait correlations (Pearson) were calculated using the cor() sfunction. Gene Ontology (GO) enrichment analyses of Biological Process terms within modules identified in WGCNA were

### Transmission electron microscopy of anemone oral tissues

Individual anemones (N = 4) were fixed by immersion in 1% v/v paraformaldehyde and 1.25% v/v glutaraldehyde in 0.1 M cacodylate buffer (pH 7.4) at an initial temperature of 35°C, followed by a 1 h incubation with constant agitation (80 rpm) at room temperature. Samples were then incubated in pos-fixative solution (2% v/v paraformaldehyde and 2.5% v/v glutaraldehyde in 0.1 M cacodylate buffer, pH 7.4) at 4°C for 24–48 h. Samples were processed and imaged at the BU Chobanian & Avedisian School of Medicine TEM core facility, using a protocol modified from previous studies^48,87,88^ as follows: tissue blocks (1–2 mm³) were rinsed in 0.1M phosphate buffer (room temperature) with 6 changes over 2 h. Tissue blocks were osmicated (1% OsO_4_ in 0.1 M phosphate buffer pH 7.4) for 2 h at room temperature, rinsed in buffer, water, and were then counterstained *en bloc* with 2% w/v uranyl acetate overnight at 4°C. Blocks were then dehydrated through a series of increasing ethanol concentrations (70–100%), infiltrated with 100% propylene oxide (Electron Microscopy Sciences, 2 x 10mins), followed by overnight infiltration in a 1:1 mixture of propylene oxide and EMbed 812 resin (Electron Microscopy Sciences) with constant agitation. The samples were then transferred to fresh 100% EMbed 812 resin for 5 h under a vacuum desiccator, and block-embedded in BEEM capsules (Ted Pella) and cured for 48 h at 60 °C. Ultrathin sections (70–90 nm) were cut using a diamond knife (Diatome) on an ultramicrotome (Leica), collected on 200-mesh copper grids (Synaptek). TEM images were acquired at 120 kV using a JEOL JEM-1400Flash transmission electron microscope (JEOL; NIH S10OD028571) equipped with an AMT NanoSprint-43M-B Mid-Mount CMOS (AMT). To quantify ER cisternal width, ten random points per image were randomly selected, and the mean width per image was calculated. Differences in ER cisternal width between treatments and symbiotic states were tested for statistical significance using a linear model and ANOVA (see Statistical Analysis).

### Quantification of Ap-NF-κB protein levels

Western blotting of Ap-NF-κB was performed as described previously^7^. First, individual anemones were homogenized using a micropestle in 2x SDS sample buffer (0.125 M Tris-HCl pH 8.0, 4.6% w/v SDS, 20% w/v glycerol, 10% v/v β-mercaptoethanol). Homogenized samples were heated at 95°C for 10 min and were then centrifuged at 13,000 rpm for 10 min at room temperature. The supernatant was transferred to a fresh 1.5 ml microcentrifuge tube. Protein samples were immediately loaded onto a 7.5% SDS-polyacrylamide gel for electrophoresis, and the proteins in the gel were subsequently transferred to a nitrocellulose membrane. The membrane was blocked for 1 h at room temperature in TBST buffer (10 mM Tris-HCl pH 7.4, 150 mM NaCl, 0.05% v/v Tween-20, containing 5% w/v non-fat dry milk). Next, the membrane was incubated with a custom anti-Ap-NF-κB antiserum^7^ (diluted 1:5000 in blocking buffer) overnight at 4°C. The membrane was then washed five times with TBST, followed by incubation with a secondary goat-anti-rabbit-HRP conjugated antiserum (1:4000, Cell Signaling Technology) for 1 h at room temperature. After an additional five TBST and two TBS washes, immunoreactive proteins were detected using the SuperSignal West Dura Extended Duration Substrate (Fisher Scientific), and Western blots were imaged using a Sapphire Biomolecular Imager. Ap-NF-κB band intensities were quantified in each lane using ImageJ, with Ponceau S staining serving as a total protein loading control. Relative Ap-NF-κB levels were calculated as the ratio of Ap-NF-κB to Ponceau S signal for each sample and were then normalized to the average of the symbiotic control (set to 1.0). Differences in Ap-NF-κB levels between treatments and symbiotic states were tested for statistical significance using a linear model and ANOVA as described above.

### Immunostaining of Ap-NF-κB

To characterize Ap-NF-κB expression in anemone tissues, immunohistochemistry was performed as described previously^7^. Briefly, anemones were fixed in 3.7% formaldehyde for 1 h at room temperature, washed with PBS for 2 min, and permeabilized in PBS containing 0.2% v/v Triton X-100 (PBSTx) for 1 h. Antigen retrieval was performed by incubating the fixed samples at 80°C for 30 min in 10 mM sodium citrate, pH 8.5, containing 0.05% v/v Tween-20. After 5 washes in PBST, samples were blocked with PBST containing 5% v/v normal goat serum and 1% w/v BSA (blocking buffer) for 1 h at room temperature. Samples were then incubated overnight with primary rabbit anti-Ap-NF-κB antiserum (1:5000 in blocking buffer) at 4°C. Samples were then incubated with Alexa Fluor 568-conjugated goat anti-rabbit IgG secondary antiserum (Invitrogen; 1:1000) for 1 h at room temperature. Following three washes in PBST for 10 min each, DNA was stained with Hoechst 33342 (Molecular Probes; 5 µM final concentration). Samples were then washed three times with PBST for 10 min each, and mounted onto glass slides. Samples were imaged on a STELLARIS confocal microscope (Leica) using channels for Alexa Flour 568 (for Ap-NF-κB), Hoechst (nuclei), and chlorophyll a (algal cell autofluorescence).

### Characterization of bacterial microbiomes

Genomic DNA was extracted from whole anemones (N = 9 per treatment per symbiotic state or 72 animals total) using an RNAqueous Total RNA Isolation Kit (Invitrogen) according to the manufacturer’s protocol. The V4 region of the bacterial 16S rRNA gene was amplified using the Hyb515F^89^ and Hyb806R^90^ primers. The PCR samples contained 1 µL of template DNA, 0.025 U ExTaq enzyme, 1x ExTaq buffer (Takara), 1 µM primers, 0.2 mM of dNTPs, and molecular biology grade water. PCR conditions were as follows: 95°C for 5 min, and 35 cycles of 1 min at 95°C, 2 min at 62°C, 2 min at 72°C, followed by 72°C for 10 min. Samples were purified using the GeneJet PCR Purification Kit (Thermo Fisher) and underwent a second PCR for dual-indexing (6 cycles using the same PCR conditions as above). Samples were pooled at approximately equimolar concentration and sequenced at Tufts University Genomics Core Facility using the MiSeq2500 platform (250 bp paired-end reads; Illumina).

Demultiplexed reads were first processed using *bbmap*^91^ to retain only reads containing the primer sequences, which were then removed using *cutadapt*^92^. Identification and quality filtering of amplicon sequence variants (ASVs) was performed using *DADA2*^93^. Taxonomic assignment was performed using *DADA2* with the Silva v.138.1 database^94^ and the National Center for Biotechnology Information nucleotide database^95^. Sequences originating from chloroplasts, mitochondria, and non-bacterial taxa were removed, and then negative control contaminants were removed using *decontam*^96^. Rarefied ASVs were trimmed with *MCMC.OTU* (5000 reads)^97^ and further processed to eliminate ASVs with low counts (<0.01%) using *vegan*^80^.

Alpha diversity indices (Shannon index, Simpson’s index, ASV richness, and evenness) were calculated using *Phyloseq*^98^. Community distance matrices based on Bray-Curtis dissimilarity were assessed using principal coordinates analysis (PCoA) with *Phyloseq* and PERMANOVAs to evaluate changes in microbiome structure and composition across treatments. All read tracking statistics are provided in table S8.

### Stable isotope analysis of carbon (δ^13^C) and nitrogen (δ^15^N)

We conducted a parallel experiment using only symbiotic anemones (n = 12 per treatment group) to quantify bulk nitrogen and carbon isotopic compositions (δ^15^N and δ^13^C) between host and algal symbionts. Following the 10-day treatment period, individual anemones were collected and homogenized as previously described^99^. Host and algal symbiont fractions were separated by centrifugation at 3,000 × *g* for 5 min at 4°C. The algal pellet and host supernatant were processed separately for isotopic analysis. All tissue samples (host and algal fractions) were freeze-dried overnight in a FreeZone benchtop freeze dryer (Labconco) until complete desiccation. Nitrogen and carbon isotopic compositions were measured using an isotope ratio mass spectrometer interfaced with a FlashSmart™ Elemental Analyzer (EA–IRMS; Thermo Fisher Scientific). Analytical precision, based on replicate measurements of internal standards and samples, was approximately ±0.2‰ and ±0.2‰ for δ^13^C and δ^15^N, respectively. Differences in δ^13^C and δ^15^N across treatments were assessed using linear models and ANOVA as described above, followed by post hoc pairwise comparisons using Tukey’s HSD test.

### Pathogen challenge experiment

To measure the effects of treatments on the suspectibility of anemones a bacterial pathogen, *Pseudomonas aeruginosa* (Pa14) was added to symbiotic anemones in various treatment groups. Symbiotic *Exaiptasia diaphana* strain CC7 was used in this additional experiment, as it exhibited the greatest fold-change increase in Ap-NF-κB protein levels in response to heat. Pathogen exposure was initiated when temperatures reached 33°C, which is when NF-κB protein and mRNA expression were both upregulated. Prior to the experiment, anemones were transferred to individual wells of six-well plates containing 10 mL of filtered seawater (FSW) and maintained in an incubator at 27°C (± 1°C) under white fluorescent light (40 µmol photons m⁻^2^ s⁻^1^) on a 12-hour light:12-hour dark cycle (light from 10:00–22:00). Experiments were conducted with n = 12 individual anemones per treatment, yielding a total of N = 60 across five treatment groups: (1) control (24°C), (2) heat, (3) heat + nitrate (5 µM), (4) pathogen only, and (5) heat + nitrate + pathogen (Fig. 7A). This design allowed us to determine the effect of pathogen exposure alone and to assess whether prior NF-κB upregulation via heat and nitrate supplementation altered survival outcomes upon pathogen challenge.

To inoculate the pathogenic bacteria, 15 µL of the bacterial suspension (∼10^7^ cells) was added directly to the individual wells of pathogen-assigned anemones. Control anemones were treated identically but received 15 µL of heat-killed *Pseudomonas aeruginosa* (Pa14). Plates were returned to the incubator and maintained at 24°C for the duration of the experiment. Survival was monitored daily for seven days post-inoculation. To compare mortality rates among treatment groups, survival analyses were conducted using the survival v.3.5-5 and survminer v.0.4.9 packages^100^. Survival probability curves were examined using a log-rank test to assess significant differences among treatment groups.

## Acknowledgments

We thank Zeba Wunderlich (Boston University [BU]), and Davies lab members for helpful discussions and feedback. We also thank the BU Supercomputing Center for computational resources. We thank Justin Scace (BU) for assistance with experimental setup and troubleshooting, Todd Blute (BU) for technical assistance with confocal imaging, and Julia Hammer Mendez (BU) for coordination of the BU Marine Program. We thank Melissa Hagy for nitrate analysis.

## Funding

National Science Foundation grant IOS-1937650 (TDG, SWD)

NSF Ocean Sciences Postdoctoral Research Fellowship award 2506815 (BHG)

BU Marine Program Warren Mcleod Summer Award, Alistair Economakis Award (JD-A)

NIH award Number S10OD028571 (MM)

Republic of China (Taiwan) Ministry of Education Scholarship, and BU Marine Program Warren

McLeod Summer Award (MC)

BU Marine Program (KT)

BU Work study (OJ)

BU Undergraduate Research Opportunities Program (KST, CC)

BU-CAMED IBIS TEM core facility, Shared Instrumentation Grant (SIG) NIH S10OD028571

## AUTHOR CONTRIBUTIONS

Conceptualization: SWD, TDG, JD-A

Methodology: JD-A, OJ, KST, CC, KT, HD, MC, FC, MP, KRM, AN, MVI, AHG, MM, RWF, TDG

Investigation: JD-A, BHG, TDG, SWD

Visualization: JD-A

Supervision: TDG, SWD

Writing—original draft: JD-A

Writing—review & editing: JD-A, OJ, KST, CC, KT, HD, MC, FC, KRM, AN, MVI, AHG, MM, RWF, TDG

